# Functional connectome reorganization relates to post-stroke motor recovery and structural disruption

**DOI:** 10.1101/2021.05.27.445834

**Authors:** Emily R Olafson, Keith W Jamison, Elizabeth M Sweeney, Hesheng Liu, Danhong Wang, Joel E Bruss, Aaron D Boes, Amy Kuceyeski

## Abstract

Motor recovery following ischemic stroke is contingent on the ability of surviving brain networks to compensate for damaged tissue. In rodent models, sensory and motor cortical representations have been shown to remap onto intact tissue around the lesion site, but remapping to more distal sites (e.g. in the contralesional hemisphere) has also been observed. Resting state functional connectivity (FC) analysis has been employed to study compensatory network adaptations in humans, but mechanisms and time course of motor recovery are not well understood. Here, we examine longitudinal FC in 23 first-episode ischemic pontine stroke patients (34-74 years old; 8 female, 15 male) and utilize a graph matching approach to identify patterns of regional functional connectivity reorganization during recovery. We quantified functional reorganization between several intervals ranging from 1 week to 6 months following stroke, and demonstrated that the areas that undergo functional reorganization most frequently are in cerebellar/subcortical networks. Brain regions with more structural connectome disruption due to the stroke also had more functional remapping over time. Finally, we show that the amount of functional reorganization between time points is correlated with the extent of motor recovery observed between those time points in the early to late subacute phases, and, furthermore, individuals with greater baseline motor impairment demonstrate more extensive early subacute functional reorganization (from one to two weeks post-stroke) and this reorganization correlates with better motor recovery at 6 months. Taken together, these results suggest that our graph matching approach can quantify recovery-relevant, whole-brain functional connectivity network reorganization after stroke.

## Introduction

Motor deficits are among the most common and disruptive symptoms of ischemic stroke. Spontaneous recovery of motor function occurs for most patients (Duncan et al., 2000), and largely depends on the ability of brain networks to functionally reorganize and compensate for lost function (Corbetta et al., 2005). As demonstrated by animal models, the functionality of damaged motor regions may be remapped to surviving tissue around the lesion (Winship and Murphy, 2009). However, brain areas distant to the lesion that have similar function and/or connectivity as the damaged area have also been shown to compensate for lost function (Winship and Murphy, 2009; Adam et al., 2020; Murata et al., 2015; Brown et al., 2009).

In humans, functional reorganization has been studied with functional magnetic resonance imaging (fMRI). Task fMRI studies have demonstrated that cortical reorganization related to motor recovery can be characterized by task-based activation of surviving ipsilesional cortex, contralesional areas, and even in regions not classically activated by motor tasks in healthy subjects (Park et al., 2011). Recently, resting-state fMRI has emerged as a means to study connectivity changes after stroke in subjects who are severely impaired and cannot perform motor tasks (Park et al., 2011). Resting-state fMRI has been used to identify recovery-related changes in functional connectivity (FC) between specific brain regions, reflecting network-level changes. However, few studies to date have attempted to capture longitudinal network-level reorganization using resting-state fMRI.

Prior studies investigating neural correlates of motor recovery have also focused almost exclusively on supratentorial strokes that impact the internal capsule and surrounding areas. Infratentorial pontine strokes may impact the corticospinal tract directly or the connections between motor cortex and the cerebellum (Lu et al., 2011), they account for roughly 7 percent of all ischemic strokes (Saia and Pantoni, 2009), and may have different mechanisms of recovery-related reorganization from those of supratentorial strokes.

Remote brain areas anatomically connected to the lesion undergo structural and functional changes through a process known as diaschisis (Carrera and Tononi, 2014). Remapping of areas anatomically connected to the lesion may be an important component of the recovery process that has not been deeply explored. In this study, we propose a novel measure to capture adaptive functional plasticity after pontine stroke, outlined below, and relate it to patterns of regional stroke-related structural connectome disruption and measures of motor recovery.

Connectivity to the rest of the brain is one aspect of a brain region’s functional role in the network. We propose that instances of functional reorganization over time, as in the case of adjacent/distal surviving tissue adopting the functional role of lost tissue, may be captured by identifying brain regions whose pattern of FC with the rest of the brain is more closely matched by a different brain region at a later date. Considering functional connectomes as a graph, the task of identifying similar nodes (gray matter regions, in this case) between two functional connectomes can be considered a graph matching problem (Conte et al., 2004). Conceptually, the process of graph matching exchanges the labels of regions in a network when doing so results in increased similarity of the two networks. When two regions exchange FC profiles, the regions are said to have been ‘remapped’. We hypothesize that graph matching, applied recently to brain networks for the first time to assess the relationship between functional and structural connectivity networks (Osmanlioǧlu et al., 2019), will allow accurate quantification of whole brain, network-level, recovery-relevant functional reorganization after stroke.

As far as we know, no work has attempted to detect connectivity network-level functional reorganization in post-stroke recovery with longitudinal MRI, much less correlate it with measures of recovery or patterns of structural connectome damage. We hypothesize that 1) brain regions with more structural damage due to the stroke will more frequently functionally reorganize, 2) more impaired subjects will have more early functional reorganization in the sub-acute stages, 3) the amount or early reorganization will be correlated with long-term recovery, and 4) the amount of functional reorganization over time will correlate with the change in motor impairment between subsequent sessions.

## Materials and methods

### Data description

The data consist of twenty-three first-episode stroke patients (34-74 years old; mean age 57 years; 8 female) with isolated pontine infarcts previously described in (Lu et al., 2011). Fourteen subjects had right brainstem infarcts, nine had left brainstem infarcts (Figure 1A). Patients were scanned five times over a period of 6 months, specifically, MRIs were obtained at 7, 14, 30, 90 and 180 days after stroke onset on a 3T TimTrio Siemens using a 12-channel phase-array head coil. Fugl-Meyer motor scores were obtained twice at each session, and later averaged and normalized to a range of 0-100 (Figure 1B). Anatomical images were acquired using a sagittal MP-RAGE three-dimensional T1-weighed sequence (TR, 1600ms; TE 2.15ms; flip angle, 9°, 1.0 mm isotropic voxels, FOV 256 x 256). Each MRI session involved between two and four runs of resting-state fMRI at 6 mins each. Subjects were instructed to stay awake with their eyes open; no other task instruction was provided. Images were acquired using the gradient-echo echo-planar pulse sequence (TR, 3000ms; TE, 30ms; flip angle, 90°, 3 mm isotropic voxels).

**Figure 1:**
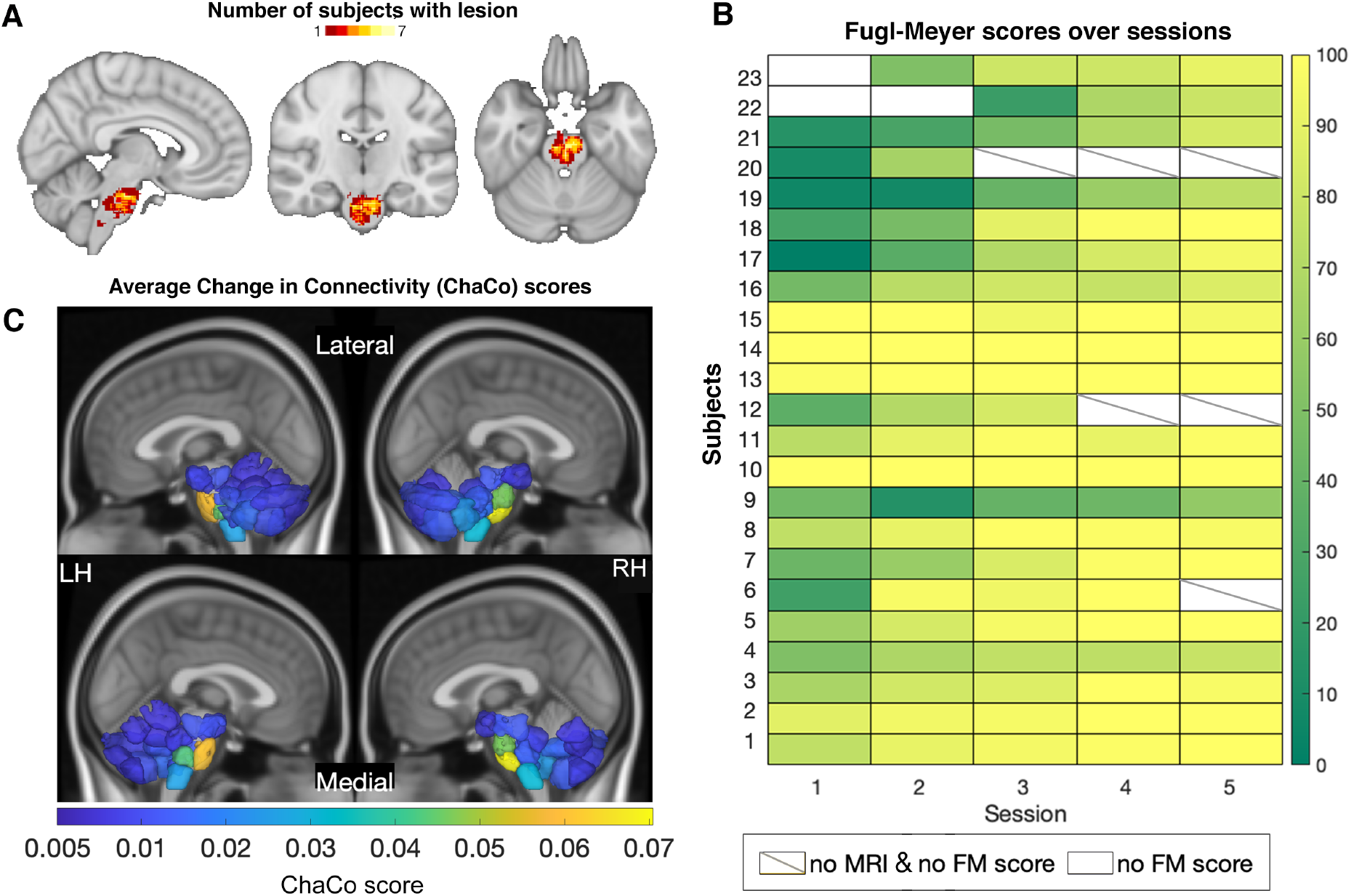
Overview of individuals’ stroke lesions, resulting structural disconnection and Fugl-Meyer score trajectories. **A.** Distribution of lesions across the brain. Colors indicate the number of subjects with a lesion in that voxel. **B.** Fugl-Meyer scores for all subjects over the five post-stroke sessions. Boxes colored white indicate missing motor scores and diagonal lines within the box indicate missing MRI data for the corresponding time point. **C.** Group average structural disconnection scores for each brain region calculated as the number of streamlines connected to each region that intersect with the lesion, normalized by the total number of streamlines connecting to that region (only displaying regions with disconnection scores >0.005). Top row of each subject inset shows a lateral view of the brain, bottom row of each inset shows a medial view.

### Structural data processing

Preprocessing of the longitudinal structural data included affine registration of each subject’s T1 scans to the baseline T1 scan, collapsing co-registered files to an average T1 and creation of a skull-stripped brain mask followed by manual editing and binarization of the hand-edited mask. The brain mask was then transformed back to each of the follow-up T1s in native space using the inverse registration acquired from the first step. This was followed by bias field correction of all the T1 scans, transformation of native-space bias field-corrected data back to baseline space, and the creation of an average bias field-corrected scan for each subject. Stroke lesion masks were hand-drawn on these transformed T1 scans by ADB and JEB. Structural normalization was performed with the CONN toolbox (Whitfield-Gabrieli and Nieto-Castanon, 2012).

### Functional data processing

Preprocessing of the longitudinal functional data was performed using the CONN toolbox (Whitfield-Gabrieli and Nieto-Castanon, 2012), including functional realignment of volumes to the baseline volume, slice timing correction for alternating acquisition, segmentation and normalization, and smoothing with a 4mm FWHM kernel. This was followed by a denoising protocol (CompCor) (Behzadi et al., 2007) which regressed out the cerebrospinal fluid and white matter signal, as well as 24 realignment parameters (added first-order derivatives and quadratic effects). Temporal band pass filtering (0.008 - 0.09Hz), despiking and global signal removal regression were also performed. The first four frames of each BOLD run were removed. Frame censoring was applied to scans with a framewise displacement threshold of 0.5 mm along with its preceding scan (Power et al., 2012). Regional time series were acquired by parcellating the scans into 268 non-overlapping brain regions using a functional atlas derived from healthy controls (Shen et al., 2013) and averaging the time course of all voxels within a given region. Voxels identified as lesioned were excluded from regional timeseries calculations. Regions were assigned to one of 8 functional networks, identified by (Finn et al., 2015) using spectral clustering in healthy subjects (Figure S1).

### Functional connectivity calculation

Functional connectivity (FC) matrices were calculated as the regularized inverse of precision matrices. Calculating FC using precision minimizes the effect of indirect connections and has been shown to result in FC that are more similar to structural connectivity (Wodeyar et al., 2020; Liégeois et al., 2020). To compute the precision FC, we first calculated the full Pearson correlation-based FC (Σ_*i*_) for each individual *i* by correlating region-pair time series. We then took the unregularized inverse of Σ_*i*_, denoted *P_i_*, and averaged them over the *i* subjects to obtain the population-level precision FC matrix *P_avg_*. We then calculated the individual precision FC matrices using Tikhonov regularization, which adds a full-rank regularization term (scaled identity) to the correlation matrix before inversion (Liégeois et al., 2020):

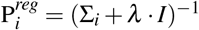

where *I* is the identity matrix and *λ* ∈ [0, 1] is the regularization parameter. The regularization parameter *λ* was chosen via a grid search to be the value that minimized the root mean squared error of the Frobenius norm of the difference in regularized subject precision matrices 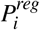 and the population-level unregularized precision matrix *P_avg_*, which was found to be *λ* = 0.73 (Fig S2). The values along the diagonal were set to 0 prior to graph matching.

### Estimated structural disconnection

Deficits from subcortical stroke may be related to functional alterations at distant sites via metabolic diaschisis (Hillis et al., 2002; Corbetta et al., 2015) or remote degeneration that spreads along the white matter connectivity network (Duering et al., 2015; Cheng et al., 2015). In order to account for the impact of lesions on the structural connectome, the extent of regional structural connectivity (SC) disruption due to the lesion was assessed for each stroke subject with the Network Modification (NeMo) Tool (Kuceyeski et al., 2013). The NeMo Tool v2 requires only an individual’s lesion mask in MNI space, which was obtained as described above, to produce an estimate of structural disconnection to each brain voxel, or to each region in a user-defined atlas. The newest version of the NeMo Tool, originally published in 2013, includes a reference database of SC from 420 unrelated individuals from the Human Connectome Project’s (HCP) 1200 release (50 percent female, aged 25-35). The NeMo Tool begins by mapping the lesion mask into this healthy database’s collection of tractography streamlines that quantify likely white matter pathways. It then identifies streamlines that pass through the lesion mask and records the gray matter voxels/regions that are at the ends of that streamline. The NeMo Tool produces the regional structural disconnection vector (ChaCo score, Change in Connectivity) that is an estimate of the percent of damaged streamlines at each voxel or region in the atlas (Figure 1C).

### Graph matching

We used a graph matching algorithm to capture FC network reorganization over time (2). Graph matching is an algorithmic process that maximizes the similarity between two networks by identifying an optimal mapping between nodes in the networks. One approach to identifying this optimal mapping is with a combinatorial optimization problem known as linear assignment.

Take two *n × n* networks *A* and *B* and a cost function *c*: 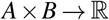 that determines the cost of assigning each node in *A* to each node in *B*. Here, our cost function is the sum of the entries in the cost matrix *C* = (*c_ij_*), whose entries *c_ij_* are defined by the Euclidean distance between row *i* in *A* and row *j* in *B*, i.e. 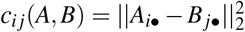. In our application, the rows *A_i•_* and *B_j•_* represent region *i* and region *j*’s FC to the rest of the brain, or FC profile, respectively. The linear assignment problem aims to construct the permutation matrix *P* = (*p_ij_*) that minimizes the sum of the elements in cost matrix, i.e. 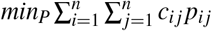. The matrix *P* is a permutation matrix with exactly one entry equal to 1 in each row and column, the rest being zero. Ones in the diagonal of *P* indicate the same node in the two networks were mapped to one another, while ones in the off-diagonal indicate a node was “remapped” to another node.

Here, we use the Hungarian algorithm to solve this minimization problem and find the corresponding optimal permutation matrix *P*. Figure 2 illustrates how the graph matching is applied to subsequent longitudinal FC networks in the same individual (either post-stroke or control) and depicts an instance of remapping in a single subject. In Figure 2A, the FC profile of brain region i (green region) at 1 week post-stroke is more closely matched by the FC profile of region j (blue region) at 2 weeks post-stroke than it is to itself at 2 weeks post-stroke. Example cost matrices for three stroke subjects (one with high amount of remapping, one with an average amount of remapping and one with a low amount of remapping) are also provided. Unsurprisingly, the lowest costs are along the diagonal (the same region mapping to itself between time points) and across left-right homologues (in the prominent super and sub diagonals).

**Figure 2:**
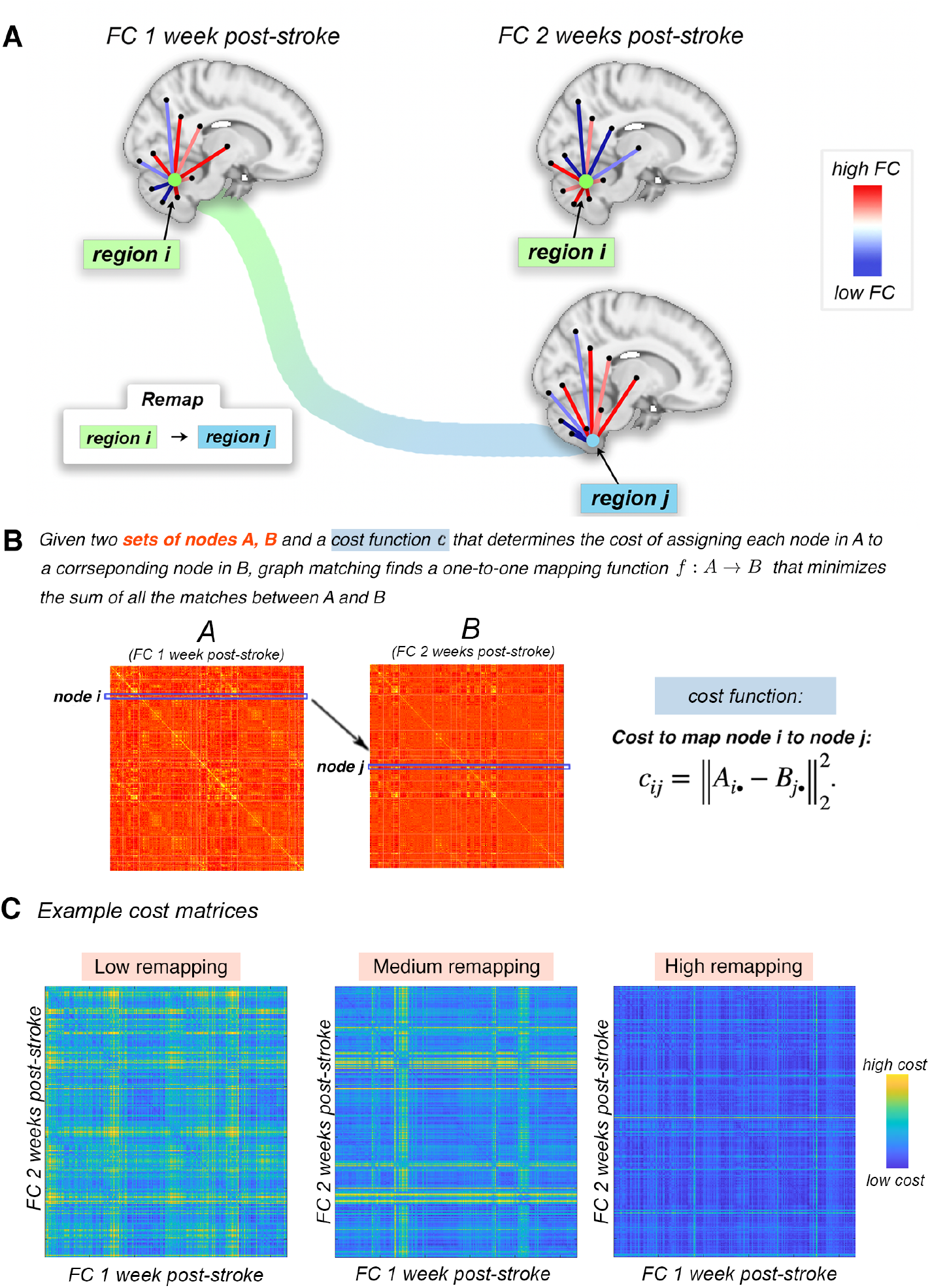
Overview of the graph matching procedure used to identify brain regions whose FC is more similar to a different region’s FC in the subsequent imaging session, i.e. regions that “remap” functional profiles. **A.** Example of a pair of regions that remap. Region *i* at 1 week post-stroke and region *j* at 2 weeks post-stroke have highly similar FC with the rest of the brain (moreso than region *i* at 1 week to region *i* at 2 weeks). The cost to remap *i* to *j* is low, and these regions would likely be remapped in the graph matching algorithm. **B.** The cost of remapping each region pair is used as input to the graph matching algorithm; the output of graph matching is the assignment of each region in one imaging session to a single corresponding region in the subsequent imaging session. If this assignment is to a different region, then it said to have “remapped”. **C.** Three examples of cost matrices of three subjects with varying amounts of remapping.

### Quantification of functional reorganization

The permutation matrices calculated for each pair of time points for each individual were used to quantify functional reorganization. To assess the spatial pattern of reorganization across the brain, we calculated each region’s ‘remap frequency’, i.e. the proportion of individuals that had that brain region remap between two subsequent time points. To assess the extent of functional reorganization for an individual, the number nodes that were remapped (sum of the off-diagonal in *P*) over time was also calculated.

### Regularization to reduce noise

We performed graph matching in two control datasets in order to identify brain regions which may remap more often due to noise or other factors. We observed remapping in controls in areas of the brainstem and cerebellum, possibly related to the low signal-to-noise ratio of these areas (Figure S3). We aimed to reduce the number of noise-related remaps by adding a regularization term to the cost matrices prior to the graph matching calculation. To do this, we added a regularization parameter to each region-pair based on the Euclidean distance between their centroids; this results in an increased cost of remapping two regions that are further apart. A Euclidean distance regularization was chosen due to the fact that remaps in control subjects were primarily between region pairs that were spatially close to each other, and because prior animal work suggests that functional remapping more often occurs in areas proximal to the lesion site. We expect that stroke-related remapping will be robust to the same amount of regularization that eliminates or minimizes remapping in controls.

Specifically, the scaled Euclidean distance matrix, whose entries *E_ij_* contain distance between centroids for two regions *i* and *j*, was added to the graph matching cost matrix *c_ij_*. Therefore, the new, penalized cost matrix is 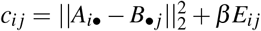, where *β* is the parameter that scales the regularization term. We varied *β* to be 0, 1e-4, 2e-4, and 3e-4 and identified the value at which remapping was minimized in controls, which was *β* = 1e-4 (Figure S4A). The few nodes that remapped in the cast-plasticity and 28andme dataset at a regularization of *β* = 1e-4 were removed in all analyses in the stroke subjects to further reduce noise.

### Statistical analyses

To test the hypothesis that brain regions more impacted by the stroke lesion would have more frequent remapping, we calculated the correlation between each region’s remapping frequency (proportion of individuals in which that region was remapped) and the sample average log-transformed ChaCo scores that quantify the regional amount of structural (white matter) connectivity disruption due to the stroke lesion. The statistical significance of the correlation was assessed with permutation testing. There are four measures of remapping frequencies that represent remapping between the four pairs of subsequent time points post-stroke (1 week - 2 weeks, 2 weeks - 1 month, 1 month - 3 months and 3 months - 6 months). Because there was a statistically significant correlation between scan length and remapping for the time point from 1 week - 2 weeks post-stroke (corr = −0.57, p(FDR)=0.01), average fMRI scan length was included as a covariate in analyses concerning remapping over time (no other time point had a significant relationship between scan length and remapping). To test the hypothesis that individuals with more remapping from one time point to another have better motor improvement, we calculated the Pearson correlation between change in Fugl-Meyer motor scores and number of remapped regions (controlling for mean fMRI scan length between sessions) from one time point to the next for all four pairs of subsequent time points. In order to leverage the repeated subject measurements, a linear mixed effects regression model was employed to determine the relationship between the amount of remapping between sessions and the change Fugl-Meyer score between sessions, including age, sex, and scan length as covariates. Let i index a subject and t index a time point. Our model is then:

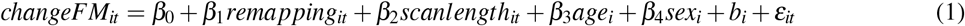

where 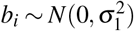 and 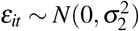. We include the random effect *b_i_* to account for correlation within a subject at different time points. To test the hypothesis that more impaired individuals would have more early remapping, Pearson’s correlation was computed between motor scores at 1 week post-stroke with the amount of remapping from 1 to 2 weeks post-stroke, controlling for the two sessions’ average fMRI scan lengths. To test if functional remapping in the early post-stroke period would be correlated with overall motor recovery, Pearson’s correlation was computed between the number of regions that remapped from 1 week to 2 weeks post-stroke and the change in Fugl-Meyer scores from 1 week to 6 months post-stroke, controlling for the two sessions’ average fMRI scan length. For the last analysis, p-values were corrected for multiple comparisons using the Benjamimi-Hochberg FDR procedure.

### Code availability

The code for to replicate this analysis is available on GitHub: https://github.com/emilyolafson/stroke-graph-matching

## Results

### Remapping in control subjects

We performed the same graph matching analysis in two separate datasets of control individuals: 1) a single female individual sampled for 30 continuous days as part of the 28andMe study (Pritschet et al., 2020) and 2) three separate individuals sampled daily for between 10 and 14 sessions as a part of a cast-induced plasticity study (Newbold et al., 2020). Several non-overlapping windows of imaging obtained 7 days apart (replicating 1 week vs 2 week comparisons) and 16 days apart (replicating 2 week vs 1 month comparisons) were extracted (see Supplementary Methods).

The **28andMe** dataset had a total of 7 non-overlapping one-week and 16-day intervals; remaps were observed in both the one-week and 16-day interval comparison (Figure S4B), with around 8% of nodes remapping without any regularization, and 0.5% of nodes remapping with a regularization level of *β* = 1e-4. The **cast-plasticity** dataset, with a total of 15 non-overlapping 1-week windows (3, 5, and 7 windows for subjects, 1, 2, and 3, respectively), also had remaps (Figure S4B), with about 6% of nodes remapping without regularization, and 0.2% of nodes remapping with a regularization level of *β* = 1e-4.

### Functional reorganization is primarily observed in the subcortical/cerebellar network

Functional remapping in stroke subjects was observed in about 18-23% of nodes (across 4 session comparisons) without regularization, and in 3-4% of nodes after imposing regularization at a level of *β* = 1e-4 (Figure S4C). Remapping occurred most often in the brainstem and cerebellum (Figure 3A,B), similar to the spatial distribution of ChaCo scores (amount of structural disconnection), which were also highest in the brainstem and cerebellum (Figure 1C). We also assessed patterns of remapping within 8 functional networks by calculating the total number of swaps across subjects that involve regions in those networks, normalized by the number of regions in each network. The most remaps occur between regions in the subcortical-cerebellum network (Figure 3C,D).

**Figure 3:**
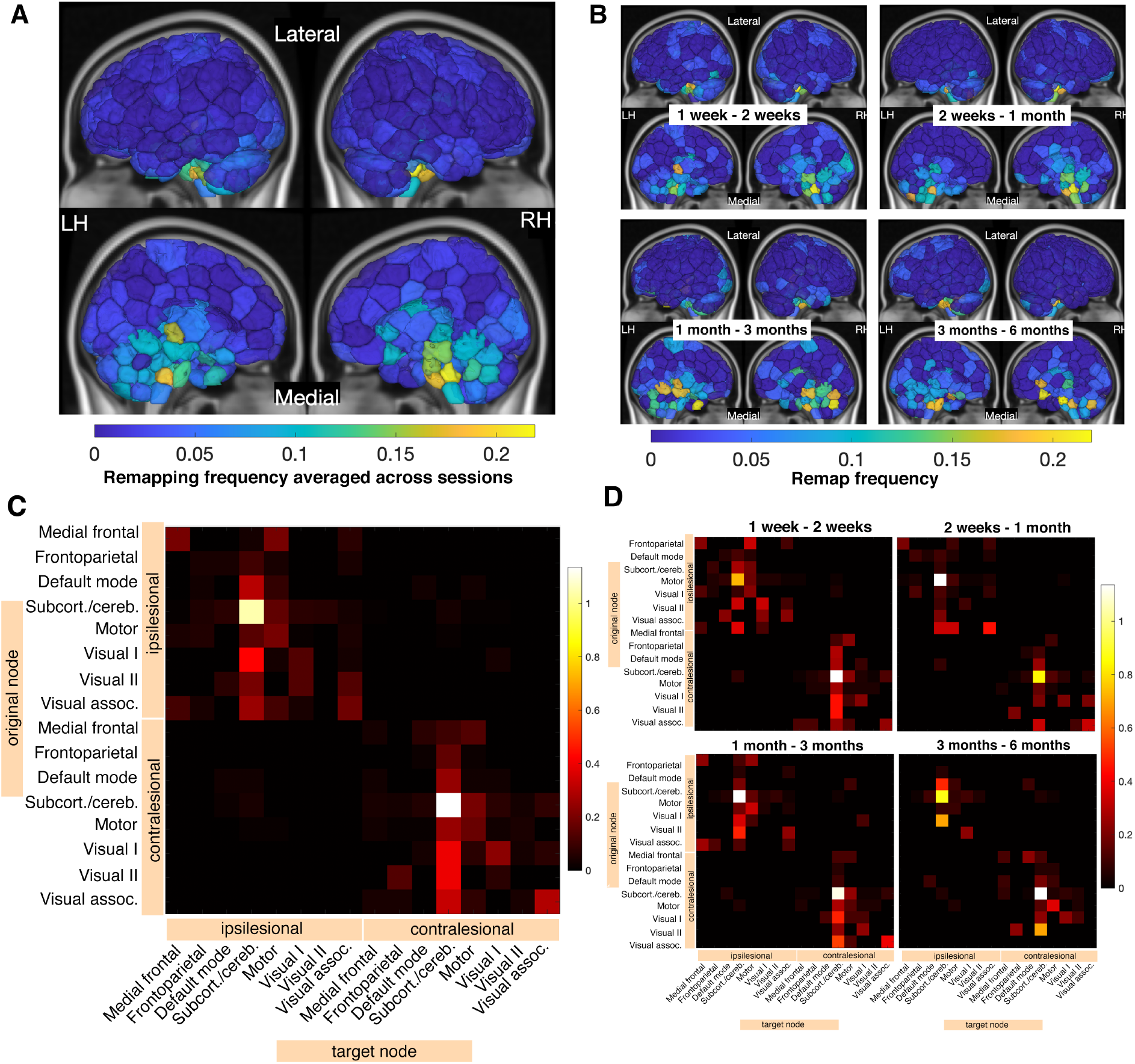
Regional functional remapping frequencies are related to mean estimated structural connectivity disruptions due to the lesion. **A.** Regional remap frequencies plotted on a glass brain, averaged across 4 inter-session comparisons. Inset figures display a lateral view (top row) and medial view (bottom row). **B.** Regional remap frequencies plotted on a glass brain, displayed separately for each inter-session comparison. **C.** Network-level sum of remaps, averaged across all 4 comparisons, and normalized by the number of regions in each network. Remaps are separated based on their position relative to the lesion (contralesional vs. ipsilesional). **D.** Network-level sum of remaps, displayed separately for each inter-session comparisons, normalized by the number of regions in each network. Ipsilesional = in the same hemisphere as the lesion, contralesional = in the opposite hemisphere as the lesion.

### Regions with greater structural connectivity network disruption have more functional reorganization over time

For each pair of subsequent time points, there was a significant, positive correlation between sample-average regional ChaCo scores and functional remapping frequency, indicating those regions with more structural connectivity disruption across the stroke subjects also had more remapping over time (Figure 4A). Furthermore, across subjects, the brain regions that remap have significantly higher ChaCo scores compared to those that do not remap (assessed with permutation testing with 10000 permutations) (Figure 4B).

**Figure 4:**
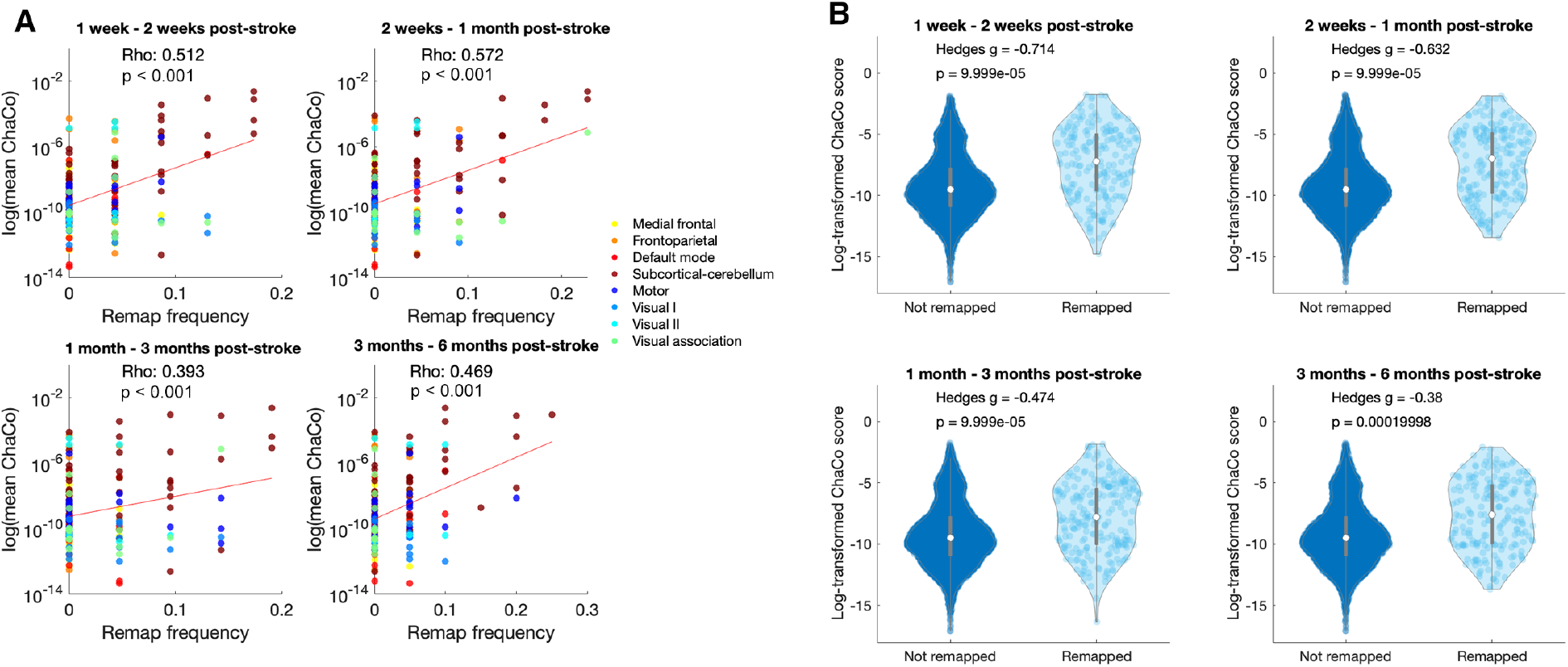
Remapping preferentially occurs in regions with greater structural connectivity disruption **A.** Correlations between average ChaCo scores and regional remap frequencies across all 23 stroke subjects. Regions are colored by network assignment. **B.** ChaCo scores of regions that remap are significantly greater than ChaCo scores of regions that do not remap.

### Functional reorganization is related to impairment and recovery

We observed a significant positive correlation between the number of early post-stroke remaps (between 1 weeks and 2 weeks post-stroke) and the 6 month improvement in motor scores, as measured by the difference in Fugl-Meyer scores at 6 months and 1 week post stroke (controlling for average scan length). This result indicates that individuals with more early functional remapping had better long-term recovery (Figure 5A). There was also a significant negative relationship between the number of early post-stroke remaps (between 1 week and 2 weeks) and baseline Fugl-Meyer scores, such that more impaired subjects had more remapping at baseline (Figure 5B). The linear mixed effects model demonstrated a statistically significant relationship between ‘remapping’ (remap) and a change in Fugel-Meyer score. For every one unit increase in ‘remapping’, there is an average increase of 0.46 units in the change in Fugl-Meyer scores, holding all other factors constant (Figure 5C)(p-value = 0.0037). The amount of recovery between subsequent sessions was significantly significant positive associated with the number of remaps between sessions for the two comparisons between 2 weeks and 1 month post-stroke (Figure 5D), but only a trend for significance existed at the 1 month to 3 months time points and there was no significant correlation for the 3 and 6 months comparison.

**Figure 5:**
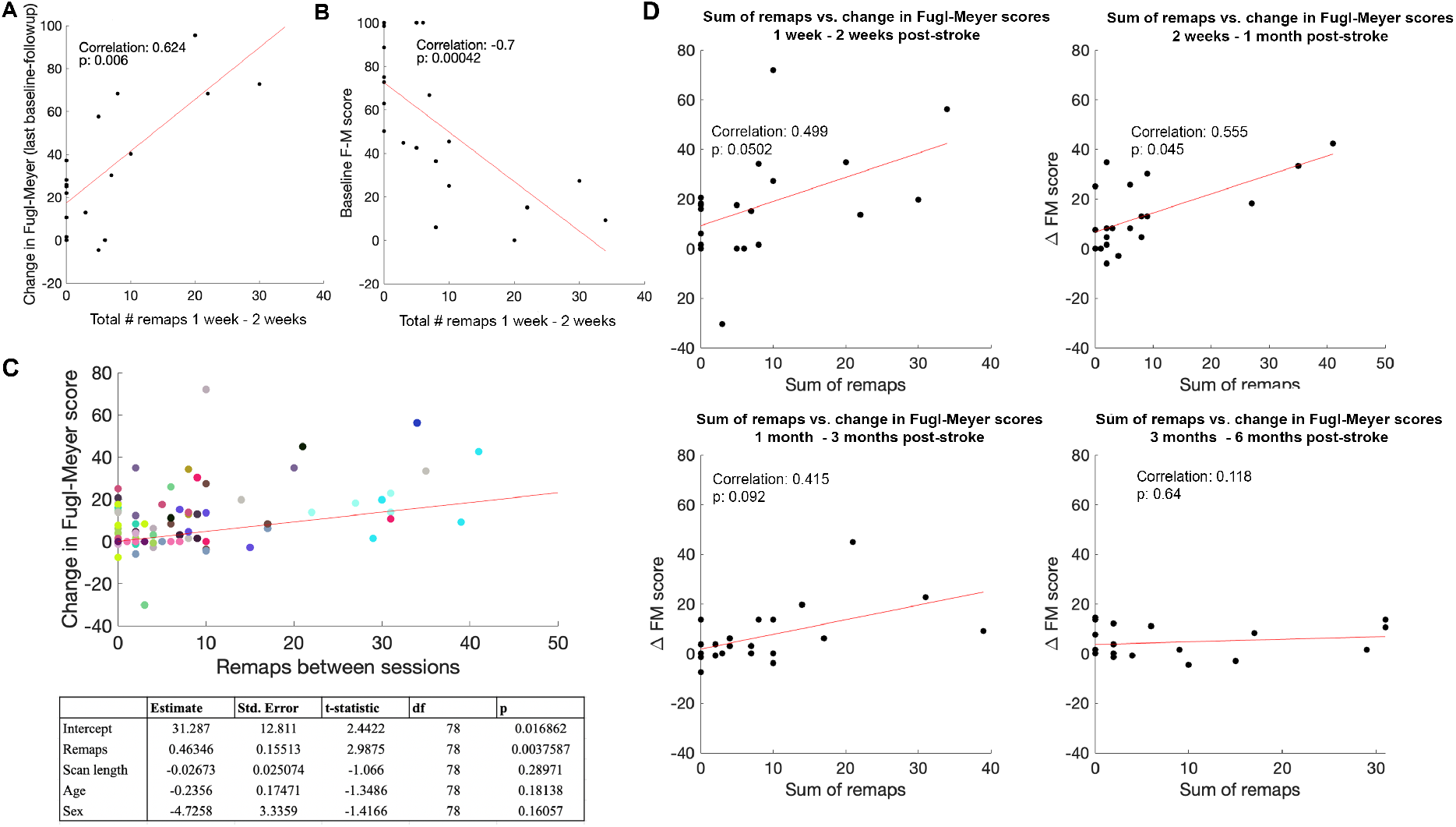
The amount of early functional remapping is related to baseline motor impairment and eventual motor recovery; functional remapping is related to change in motor recovery between subsequent time points. **A.** Pearson correlation between individuals’ total number of early remaps (between 1 week FC and 2 week FC) and their change in Fugl-Meyer scores between 1 week and 6 months post-stroke (Spearman correlation: 0.549, p(FDR)=0.022). **B.** Pearson correlation between individuals’ total number of early remaps (between 1 week FC and 2 week FC) and 1 week Fugl-Meyer scores, controlling for subjects’ average scan lengths between 1 week post-stroke and session 2 (Spearman correlation: −0.654, p(FDR) = 0.002). **C.** Results from linear mixed effects analysis; colors indicate different individuals’ longitudinal time points. **D.** Pearson correlation between the total remaps observed between each pair of time points and the change in Fugl-Meyer scores between time points, controlling for average fMRI scan length. P-values are corrected for multiple comparisons (Spearman correlations: Corr. 1 week-2 weeks = 0.456, p(FDR)=0.0575, Corr. 2 weeks - 1 month = 0.566, p(FDR)=0.03, Corr. 1 month - 3 months = 0.533, p(FDR)=0.03, Corr. 3 months - 6 months = −0.09, p(FDR)=0.714)

### Robustness of results

The impact of lesions introducing noise into the BOLD signal of underlying gray matter is likely not driving remapping, as the maximum overlap of the lesions with each ROI is no more than 30 percent (Figure S5) and, importantly, lesioned voxels were excluded from FC calculations. We also found that the number of remaps was not related to differences in in-scanner motion between scans, as measured by framewise displacement (Fig S6). Additionally, there is one free parameter in the graph matching, which was the amount of regularization (value of *β*). The main results we present here based on a value that reduced the amount of remapping in controls to a level of <1% (see Figure S4). However, we also replicated the main results across an increasing value of *β* to show robustness of the findings to this parameter choice (Figure S8, S9, S10).

## Discussion

In this paper, we proposed a measure of functional connectome reorganization based on graph matching, and evaluated its relationship to structural disconnection and motor impairment/recovery in a set of 23 individuals with pontine stroke. We observed instances of functional reorganization over the 1 week to 6 months post-stroke period, and demonstrated that the areas that undergo functional reorganization most frequently are in cerebellar/subcortical networks. Furthermore, regions more impacted by stroke via disruption to their structural connections had more functional remapping over time. Finally, we show that functional reorganization one week post-stroke is highly related to both baseline impairment and the extent of motor recovery at 6 months, and, finally, that the extent of functional reorganization between 2 weeks and 1 month post-stroke is correlated with the extent of motor recovery observed in this same subacute time period.

Motor recovery following stroke is supported by active functional and structural remodelling in the area bordering the stroke, including increases in excitability, increases in dendritic spine turnover, and the formation of new axonal projections (Murphy and Corbett, 2009). This remodelling can entail surviving cortical areas remapping functionally to compensate for post-stroke impairment (Brown et al., 2009). The dynamic reorganization of resting-state functional connectivity, observed in this stroke cohort in cerebellum and subcortical areas, may reflect longitudinal compensatory changes in representations of motor functions. Interestingly, we showed correlations between amount of functional remapping and amount of motor improvement but only in the period of time in which most post-stroke recovery occurs (within 3 months after stroke) (Lee et al., 2015). However, resting-state functional connectivity, though related to task-based responses, may not be fully representative of brain activation patterns underlying specific behaviors, and has shown to be constrained by the structural connectome (Kuceyeski et al., 2019; Honey et al., 2009). Thus, it is possible that the remapping observed is not a functional remapping per se (in the sense of remapping a region’s role in a specific task), but a shift in balance of a brain area’s functional connectivity profile at rest, which could reflect task-based functional compensations, changes in network topology, and/or underlying structural remodelling.

We also show that more remapping between sessions is associated with better motor recovery between those same sessions, and that more remapping in the early phase of stroke is related to both amount of baseline impairment and the degree of motor recovery at 6 months. Functional reorganization measured with this remapping technique may be interpreted as reflecting a compensatory, beneficial process that is proportional to the extent of motor recovery. On the other hand, this approach may also be capturing phenomena that occurs during the first 6 months after stroke that are unrelated to recovered functionality, but that correlate with measures of recovery nonetheless. For instance, stroke-related increases in noise in the subcortical/cerebellar may result in more remapping which could be the result of a random or pathological process concomitant with a recovery mechanism. Thus, more densely-sampled fMRI studies should be performed to identify other elements of this reorganization process.

The most remaps were observed in the subcortical/cerebellum network, suggesting that remapping is likely reflecting a process of functional reorganization that is spatially constrained (Figure 3). This network-level reorganization is consistent with prior task-based studies showing remapping of motor-based activations to premotor and homologous areas, and with studies of resting-state FC that demonstrate specific spatial patterns of FC changes after stroke, like changes in resting state FC within motor networks (Zhang et al., 2016), and contralesional functional connectivity changes that are concentrated in a small set of brain regions (Yourganov et al., 2021).

We now understand that deficits arising from stroke are not only related to the damage inflicted at the stroke core, but also to remote cortical areas structurally connected to the lesion, due to retro- and anterograde degeneration (Guggisberg et al., 2019). For instance, several studies have shown that there are local reductions in cortical thickness in areas directly connected to subcortical lesions (Duering et al., 2015; Cheng et al., 2015). Here, we show that the amount of structural disconnection from a stroke lesion is associated with greater functional reorganization after stroke, suggesting that indirect damage to region’s structural connections may also trigger its functional remapping.

Plasticity is frequently observed in the area bordering the stroke (Brown et al., 2009), where tissue may have a similar function to the area damaged by stroke. To incorporate the a priori hypothesis that recovery-related remapping would be observed between adjacent regions, we added a penalty to the graph matching algorithm proportional to the distance between region-pairs. This penalty term did not substantially alter our results, suggesting that the relationships we observed between remapping and recovery were driven, in part, by remapping between areas that were closer together. Long-range functional reorganization has been observed in animal models, particularly when the lesion is large. However, most studies of ‘smaller’ strokes in rodent models resemble the relative size of a survivable human stroke (Murphy and Corbett, 2009), so we assume that the stroke subjects we observed had relatively smaller strokes that would more likely have remapping between adjacent regions.

### Limitations

There are several limitations to this study. The first is that the impact of noise cannot be completely accounted for due to a lack of sufficient controls with the same acquisition and processing methods. It is possible that the nodes that remap more frequently are in brain areas that have lower SNR and that remaps observed as indicative of noisy signal. Indeed, the SNR of HCP subjects is lowest in medial structures such as the thalamus, cortical midsurface, and most consequentially to this study, the anterior cerebellum (see S3).However, the SNR of the subcortical/cerebellum network was not sufficient to cause substantial remapping in the longitudinal data in healthy individuals after regularization. Finally, the control analyses used data from two different studies with varying scanners and processing pipelines but minimal remapping was observed, supporting the hypothesis that fMRI noise is not driving remapping.

### Conclusions

In this study we proposed a measure of functional connectome reorganization, called remapping, and applied it to longitudinal resting-state fMRI data in a cohort of pontine stroke patients. Remapping was observed in all stroke subjects and was correlated with recovery over the early to late subacute phases. Areas impacted by the lesion through structural disconnection were more likely to remap than those areas not impacted by the lesion. Two independent analyses showed that little to no remapping is observed in healthy individuals. This work expands our understanding of functional processes related to recovery after stroke. If we can identify subjects who have more potential for functional reorganization, or can devise therapeutics to boost this remapping mechanism, we may be able to improve patient outcomes after stroke.

## Acknowledgements

We thank Dr. Daniel Tranel, Dr. Zhou Fan, and Chang Su for their valuable and constructive suggestions during the development of this research work.

## Supplementary Methods

### Supplementary control analyses

#### 28andMe data

The 28andMe dataset (Pritschet et al., 2020 consists of densely sampled anatomical and functional MRI on a female participant (23 y.o.) who underwent imaging for 30 consecutive days. fMRI preprocessing is detailed in (Pritschet et al., 2020; details do not deviate significantly from the processing described in this paper. Because the TR for the resting state fMRI obtained for this subject was much lower than the TR of stroke subjects (720ms vs 3000ms, respectively), the 28andMe data was downsampled to an equivalent TR. Initial scans contained 820 frames at a TR of 720ms, equating to roughly 10 minutes of acquisition. Resampling to 1/4th of the frequency was performed using MATLAB’s *resample()* function which performs downsampling as well as filtering to account for aliasing, which reduced the number of frames to 206 for each of the 30 scans. GSR was then performed on the regional timeseries data.The 28andMe analyses were completed using functional connectivity derived from the linear correlation between region time series. Seven sets of scans were extracted (day 1, 8, 24; day 2, 9, 25; day 3, 10, 26; day 4, 11, 27; day 5, 12, 28; day 6, 13, 29; and 1 week, 14, 30), with intervals between the days selected to replicate the interval between scanning in the stroke subject’s first three visits (7, 14 and 30 days post-stroke). Graph matching was applied to each of the 7 sets of scans for both pairs of subsequent time points within the set, and the resulting remapping frequencies calculated over the 14 total sets of FCs.

#### Cast-induced plasticity data

Each of three adult subjects were scanned daily for a cast-induced plasticity study by Newbold et al. (Newbold et al., 2020), two were males (35 y.o. and 27 y.o.) and one was a 25 y.o. female. Subjects underwent structural and functional MRI scanning for 10-14 days before their dominant arm was covered in a medical cast, and were scanned for a period of time during the cast and after the cast was removed. Because significant resting-state functional connectivity changes were observed during the cast period, we did not use this data, nor did we use the data following the cast removal because a different scanner was used. Participants were scanned for 30 minutes using a 3T Siemens Trio MRI scanner; we only used 14 minutes of scan data (first 380 TRs) in order to match the mean scan length of our stroke subjects. BOLD data were acquired at a spatial resolution of 4mm, single-band, with a TR of 2.2s. Subject 1 had 10 pre-cast scans, subject 2 had 12, and subject 3 had 14. These subjects were used to investigate remapping in controls in the 1-week window that replicates the difference between 1 and 2 weeks post-stroke in the stroke subjects. For each subject, we obtained several non-overlapping windows. Specifically for subject 1, we compared: day 1 and day 8, day 2, 9, and day 3, 10. For subject 2, we compared: day 1, 8, day 2, 9, day 3, 10, day 4, 11, day 5, 12. For subject 3, we compared: day 1, 8, day 2, 9, day 3, 10, day 4, 11, day 5, 12, day 6, 13, 1 week, 14. Raw data was downloaded from Datalad and processed with the same pipeline as the stroke subjects. Graph matching was applied to the resulting scans.

### Supplementary Figures

**Figure S1:**
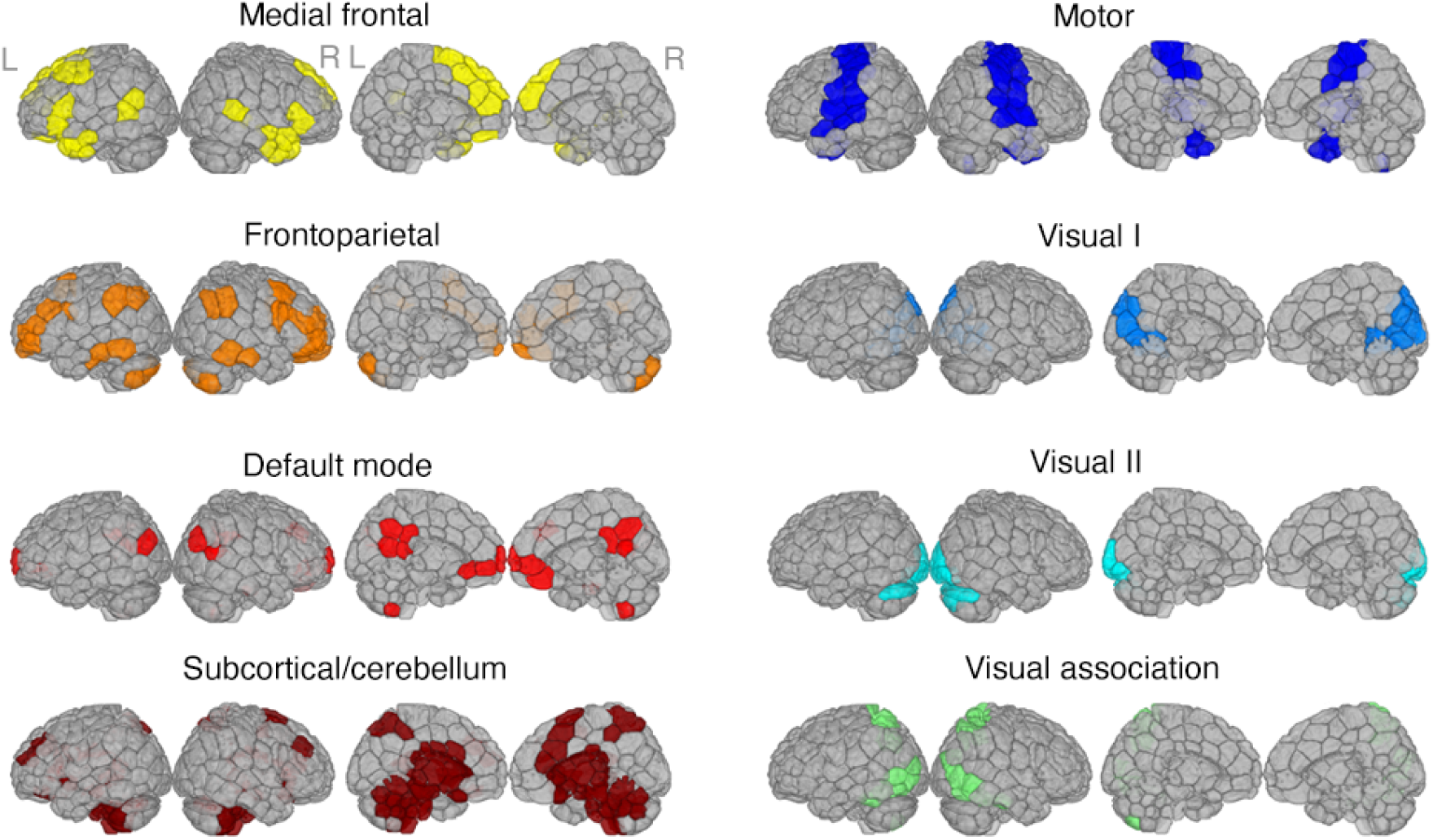
8 functional networks identified by (Finn et al., 2015) by clustering healthy functional connectivity matrices.

**Figure S2:**
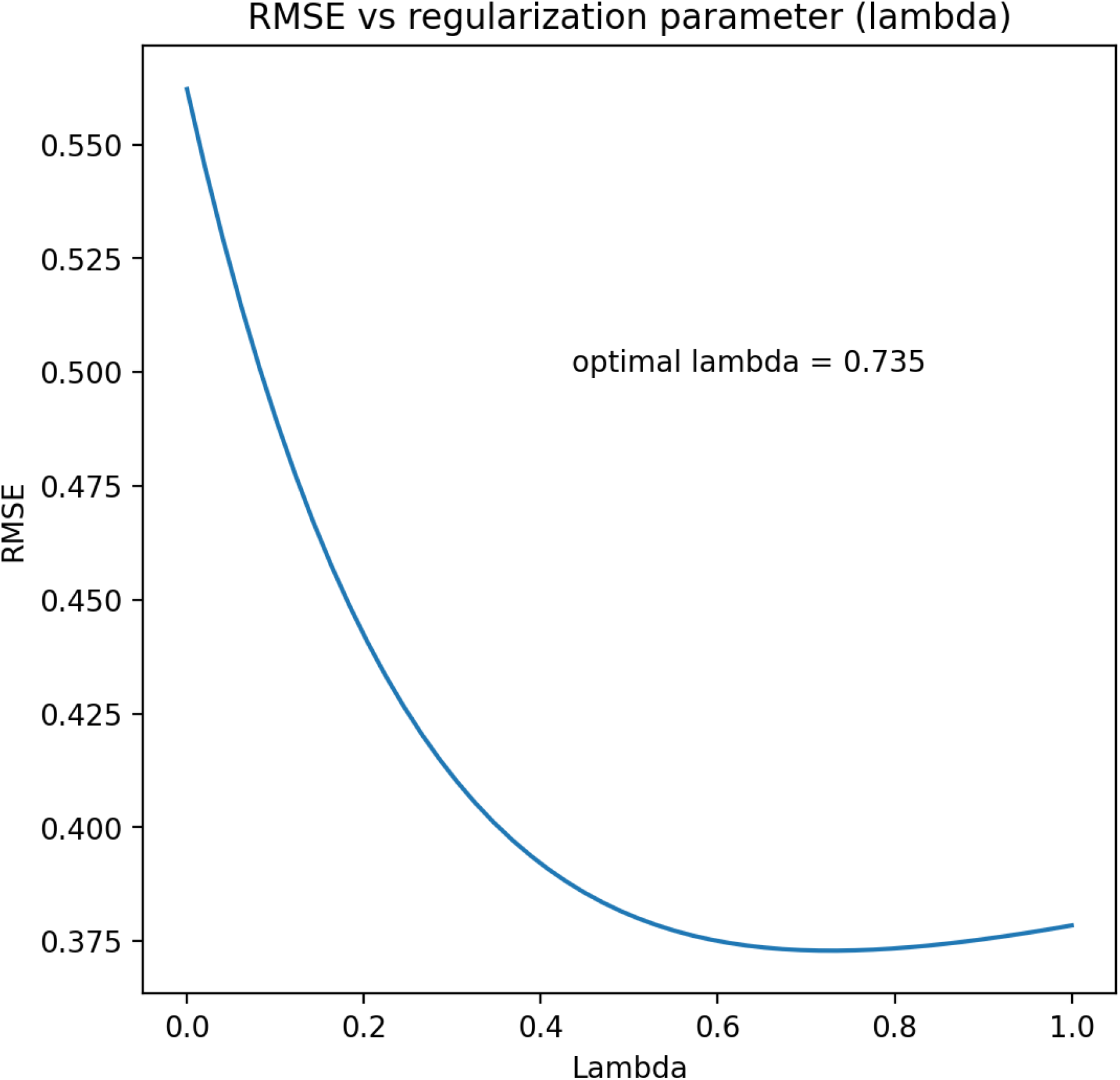
Lambda parameter that minimizes the error of precision matrices

**Figure S3:**
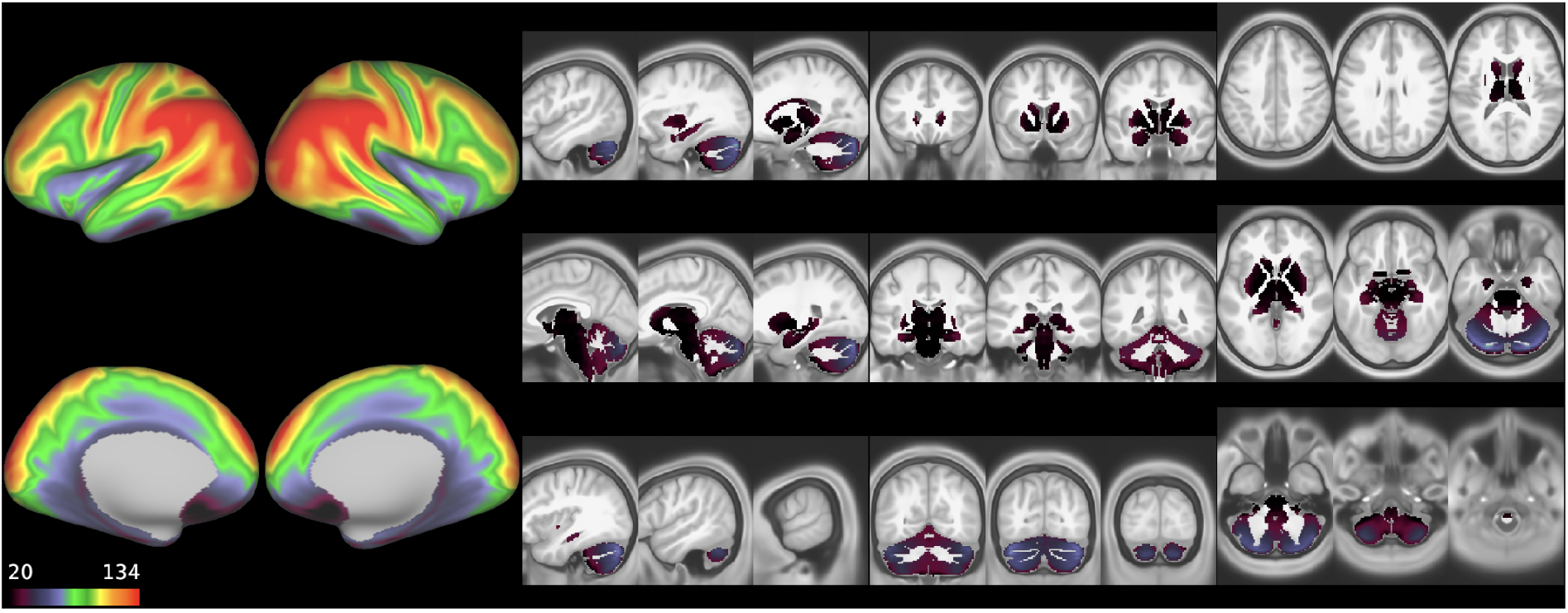
SNR calculations on the test-retest HCP BOLD data: temporal signal-to-noise ratio: TSNR=MEAN/stdev(non-artifact thermal noise)

**Figure S4:**
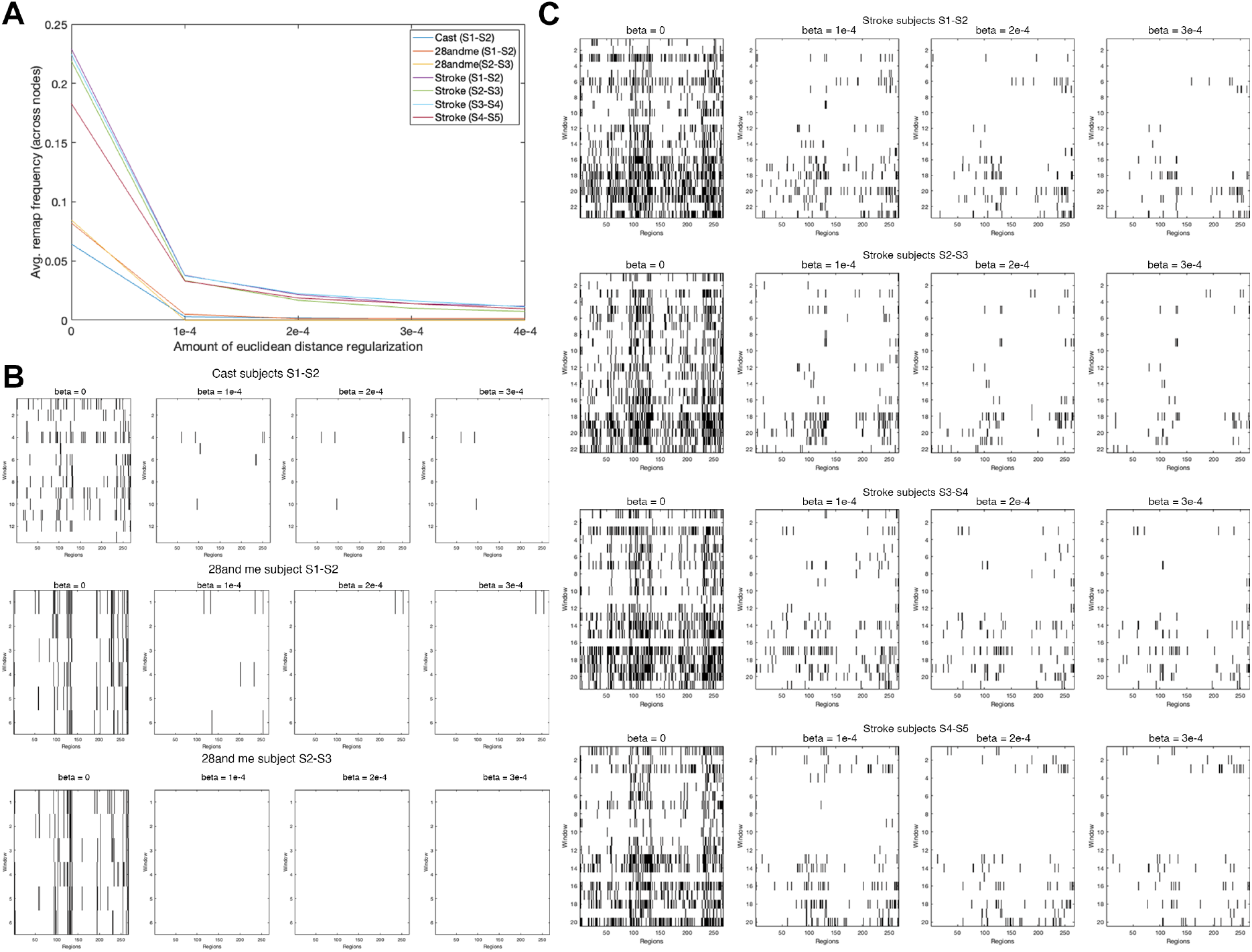
A. Average remap frequency (average across nodes) of the control datasets (cast and 28andme) relative to stroke subjects, with varying levels of regularization. The amount of regularization that minimized remapping in controls (1e-4) was used to regularize stroke subjects. B. Remapping of cast-induced plasticity controls (top row) and of 28andme subject (bottom 2 rows) with varying levels of euclidean distance regularization. C. Remapping of stroke subjects with varying regularizaiton.

**Figure S5:**
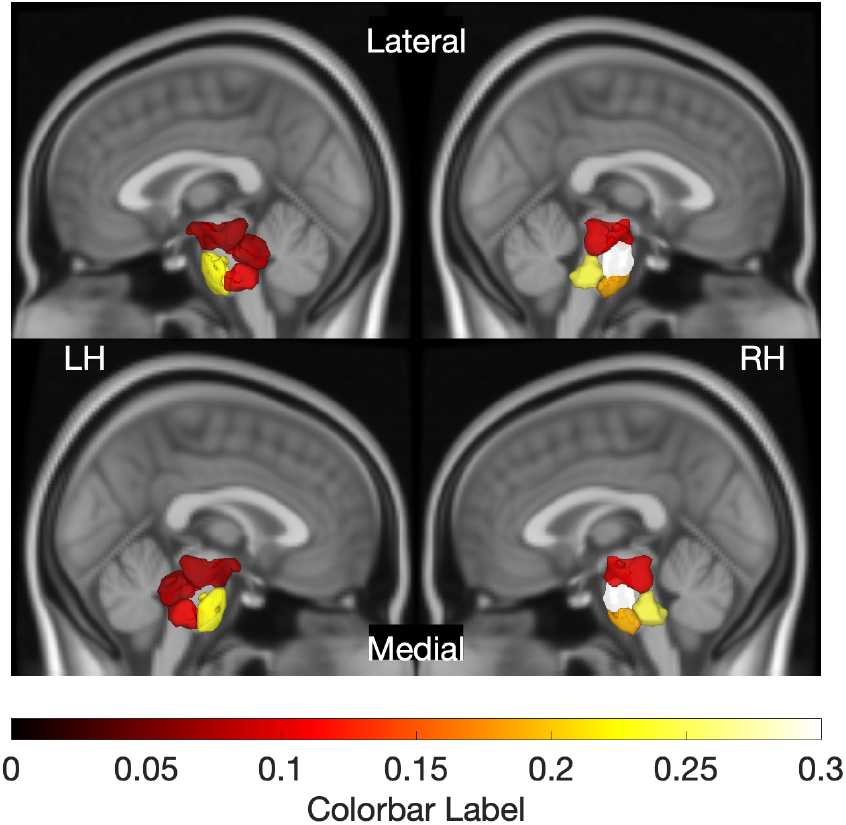
Average overlap between each gray matter regions and lesions; calculated for each subject as the total volume of the lesion inside each region divided by the total region volume. Displaying only regions with an average overlap with the lesion above zero.

**Figure S6:**
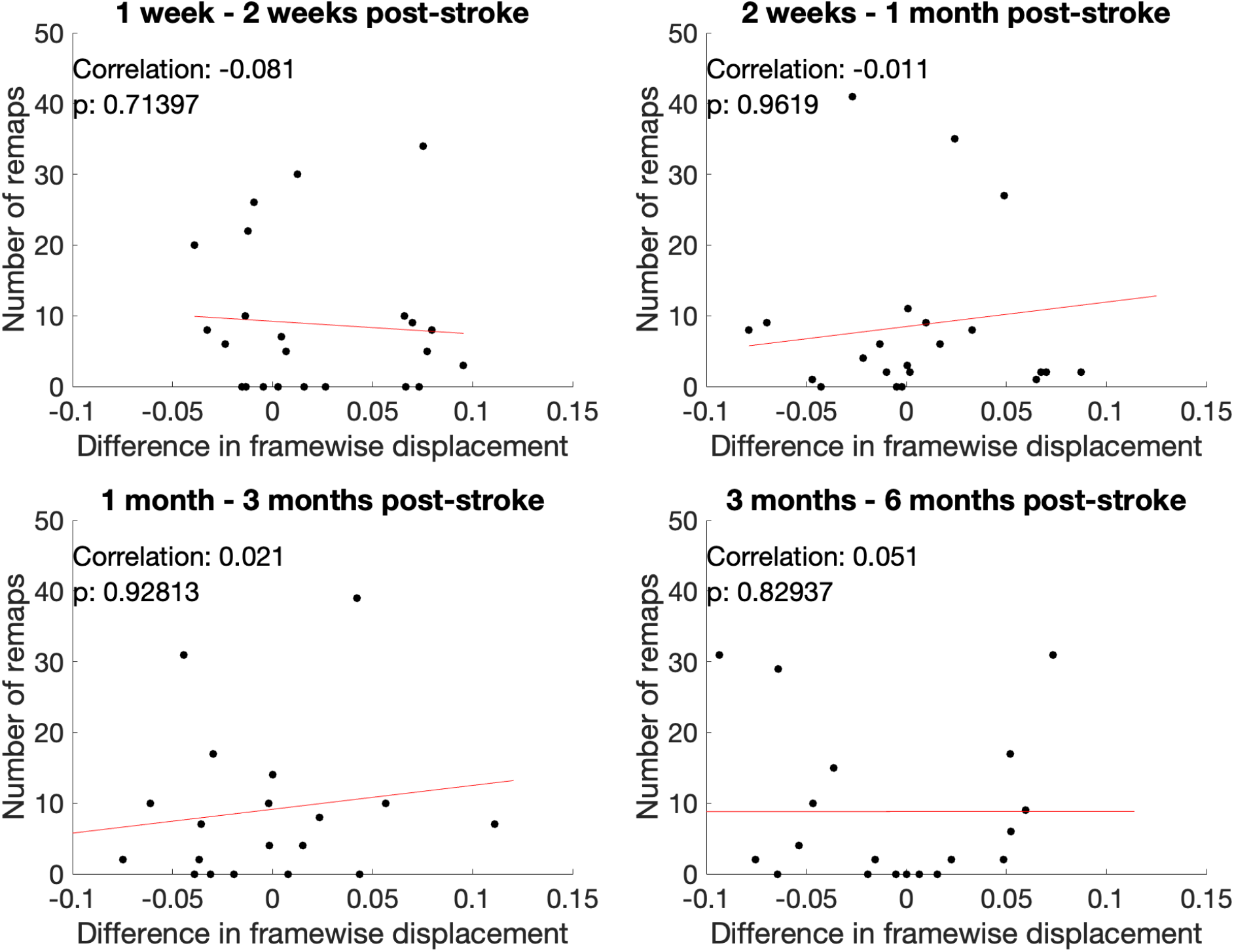
Average framewise displacement between scans does not correlate with the sum of remaps observed between scans.

**Figure S7:**
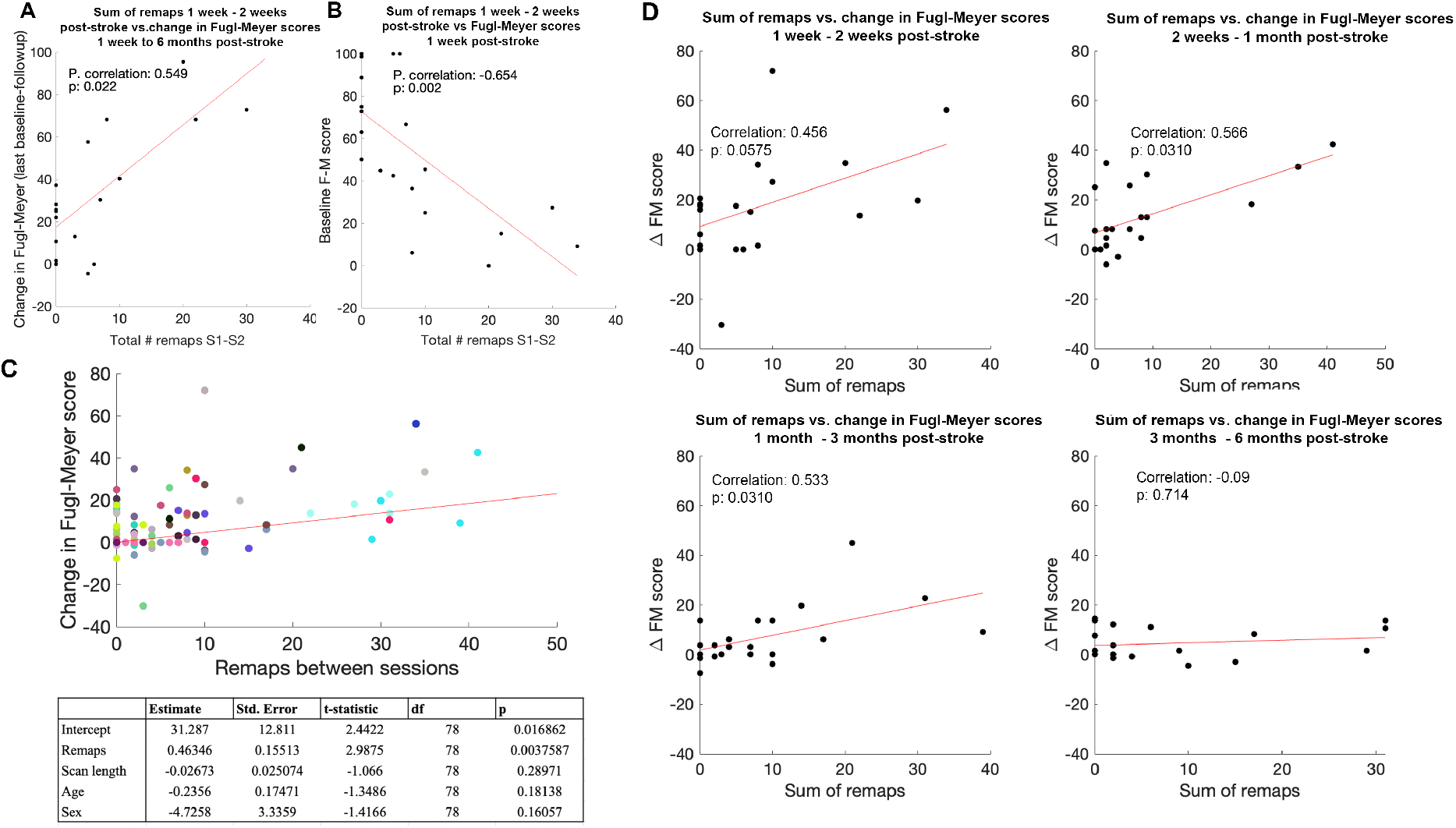
The amount of early functional remapping is related to baseline motor impairment and eventual motor recovery; functional remapping is related to change in motor recovery between subsequent time points. **A.** Spearman correlation between individuals’ total number of early remaps (between 1 week FC and 2 week FC) and their change in Fugl-Meyer scores between 1 week and 6 months post-stroke. **B.** Spearman correlation between individuals’ total number of early remaps (between 1 week FC and 2 week FC) and 1 week Fugl-Meyer scores, controlling for subjects’ average scan lengths between 1 week post-stroke and session 2. **C.** Results from linear mixed effects analysis. **D.** Spearman correlation between the total remaps observed between each pair of time points and the change in Fugl-Meyer scores between time points, controlling for average fMRI scan length. P-values are corrected for multiple comparisons.

**Figure S8:**
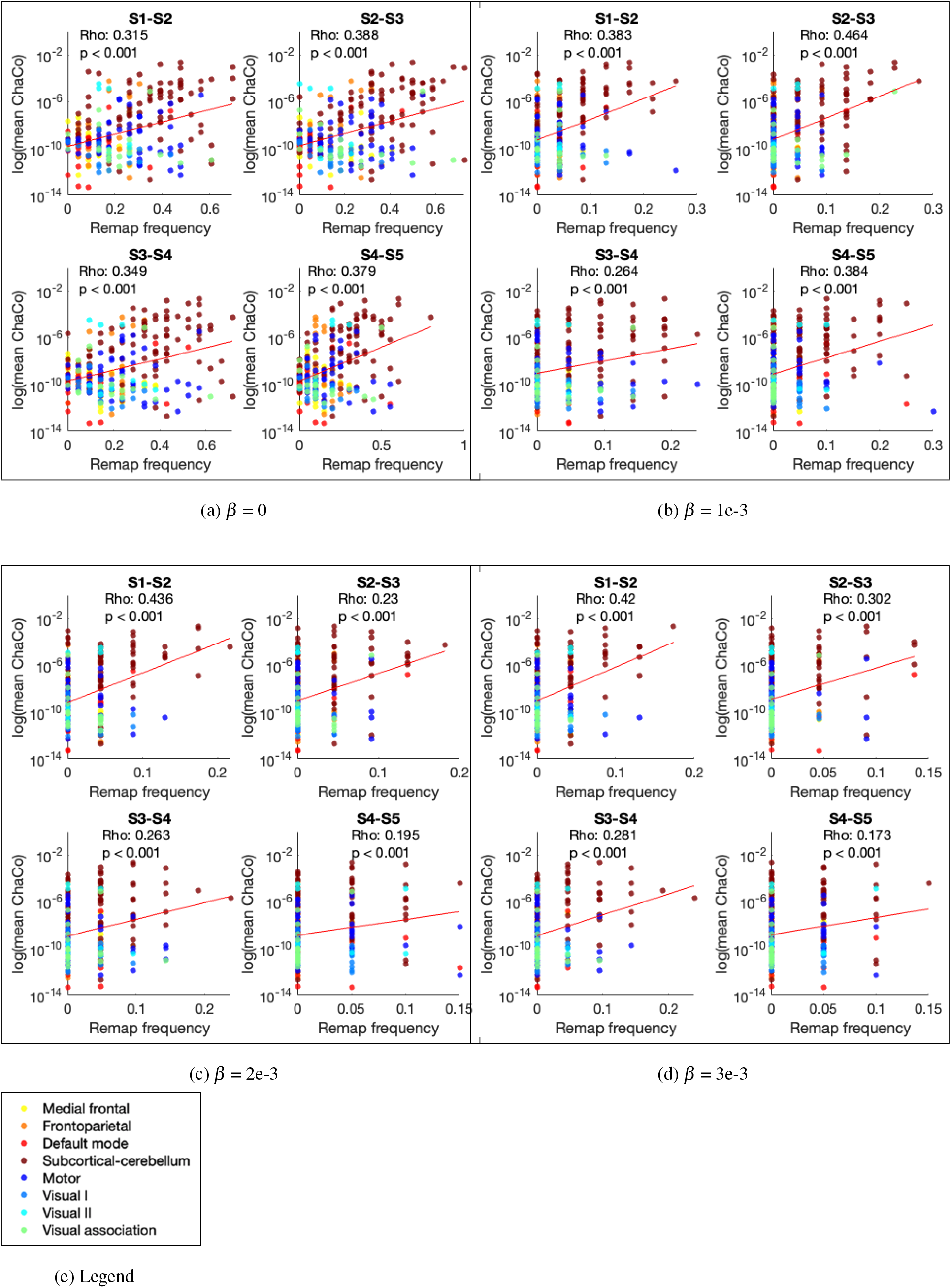
Varying *β* does not impact the significance of the correlation between regional number of remaps and ChaCo scores. P-values are corrected for multiple comparisons.

**Figure S9:**
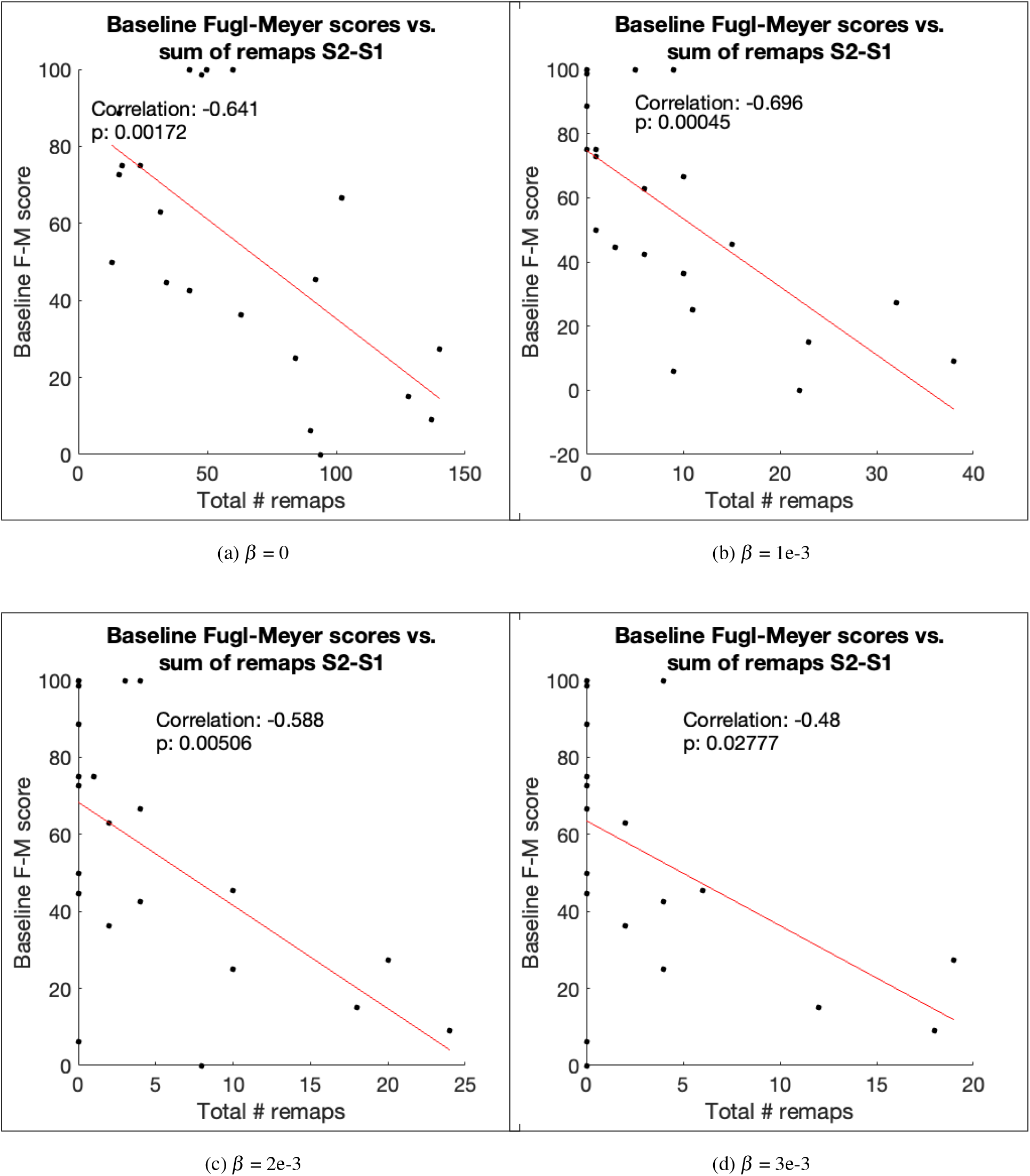
Varying *β* does not change the significance of the relationship between total number of remaps between 1 and 2 weeks post-stroke and a) baseline Fugl-Meyer scores or b) 6-month follow-up Fugl-Meyer scores.

**Figure S10:**
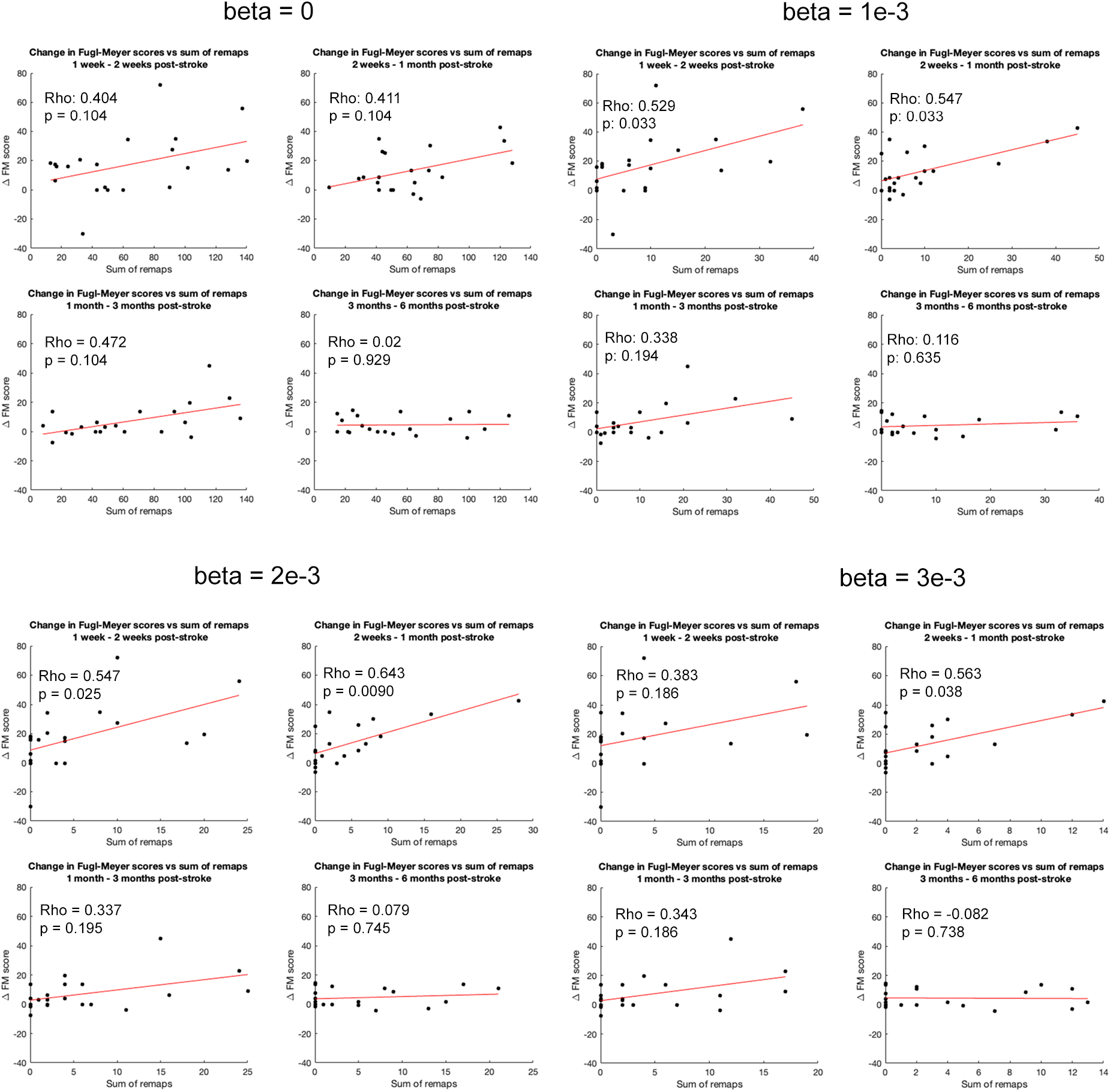
Varying *β* generally does not alter the relationships observed between session-specific changes in Fugl-Meyer motor scores and the number of regions that functionally remap between sessions. Corrected p-values are reported.

